# SARS-CoV-2 variant B.1.617 is resistant to Bamlanivimab and evades antibodies induced by infection and vaccination

**DOI:** 10.1101/2021.05.04.442663

**Authors:** Markus Hoffmann, Heike Hofmann-Winkler, Nadine Krüger, Amy Kempf, Inga Nehlmeier, Luise Graichen, Anzhalika Sidarovich, Anna-Sophie Moldenhauer, Martin S. Winkler, Sebastian Schulz, Hans-Martin Jäck, Metodi V. Stankov, Georg M. N. Behrens, Stefan Pöhlmann

## Abstract

The emergence of SARS-CoV-2 variants threatens efforts to contain the COVID-19 pandemic. The number of COVID-19 cases and deaths in India has risen steeply in recent weeks and a novel SARS-CoV-2 variant, B.1.617, is believed to be responsible for many of these cases. The spike protein of B.1.617 harbors two mutations in the receptor binding domain, which interacts with the ACE2 receptor and constitutes the main target of neutralizing antibodies. Therefore, we analyzed whether B.1.617 is more adept in entering cells and/or evades antibody responses. B.1.617 entered two out of eight cell lines tested with slightly increased efficiency and was blocked by entry inhibitors. In contrast, B.1.617 was resistant against Bamlanivimab, an antibody used for COVID-19 treatment. Finally, B.1.617 evaded antibodies induced by infection or vaccination, although with moderate efficiency. Collectively, our study reveals that antibody evasion of B.1.617 may contribute to the rapid spread of this variant.

## INTRODUCTION

The severe acute respiratory syndrome coronavirus 2 (SARS-CoV-2) is the causative agent of the devastating coronavirus disease 2019 (COVID-19) pandemic, which more than one year after its emergence is associated with record numbers in cases and deaths (W.H.O., 2021). Effective antivirals are largely lacking, although recombinant antibodies targeting the viral spike protein (S) can significantly reduce viral load und have received emergency use authorization (EUA) (Chen et al., 2021a; Gottlieb et al., 2021). Drugs that target the dysregulated cytokine responses characteristic for COVID-19 are available but their clinical benefit is oversee-able (Tomazini et al., 2020; W.H.O. REACT Working Group et al., 2020). While progress in treatment development is moderate, mRNA- and vector-based vaccines are available that provide efficient protection against disease (Baden et al., 2021; Polack et al., 2020). As a consequence, vaccination is viewed as the key instrument in combating and ultimately ending the COVID-19 pandemic.

Strategies to fight the COVID-19 pandemic either by vaccines or non-pharmaceutical interventions have been threatened by the emergence of SARS-CoV-2 variants of concern (VOC). These variants harbor mutations that confer increased transmissibility or immune evasion (Plante et al., 2021). Studies of mutations present in VOC have mainly focused on the viral S protein. The S protein is incorporated into the viral membrane and facilitates viral entry into target cells. For this, the surface unit, S1, of the S protein first binds to the cellular receptor ACE2 (Hoffmann et al., 2020; Zhou et al., 2020) via its receptor binding domain (RBD). Subsequently, the S protein is activated by TMPRSS2 or related cellular proteases (Hoffmann et al., 2021b; Hoffmann et al., 2020) and the transmembrane unit, S2, of the S protein facilitates fusion of the viral and a cellular membrane, allowing delivery of the viral genome into the host cell. These processes are essential for SARS-CoV-2 infection and are targeted by drugs and neutralizing antibodies.

The prototypic VOC with increased fitness is variant B.1.1.7, which emerged in the United Kingdom and is now spreading in many countries. B.1.1.7 replicates to higher levels in patients and is more efficiently transmitted between humans as compared to the previously circulating viruses (Frampton et al., 2021; Graham et al., 2021; Leung et al., 2021). The increased transmissibility might be linked to mutation N501Y in the RBD that might increase binding to ACE2 (Ali et al., 2021; Luan et al., 2021). However, the exact mechanisms underlying more robust transmission of B.1.1.7 remain to be elucidated. In contrast, differences in antibody-mediated neutralization between previously circulating viruses and variant B.1.1.7 are minor, with B.1.1.7 being slightly less sensitive to neutralization (Chen et al., 2021b; Collier et al., 2021; Hoffmann et al., 2021a; Kuzmina et al., 2021; Muik et al., 2021; Planas et al., 2021; Shen et al., 2021; Supasa et al., 2021; Wang et al., 2021a; Xie et al., 2021). In sum, B.1.1.7 shows increased fitness and will outcompete previously circulating viruses in an immunologically naïve population.

In populations with a high percentage of individuals with pre-existing immune responses against SARS-CoV-2, viral variants that can evade immune control have a selective advantage. Variant B.1.351 that became dominant in South Africa (Tegally et al., 2021) and variant P.1 that became dominant in Brazil (Faria et al., 2021) are such variants (Chen et al., 2021b; Dejnirattisai et al., 2021; Edara et al., 2021; Garcia-Beltran et al., 2021; Hoffmann et al., 2021a; Kuzmina et al., 2021; Planas et al., 2021; Wang et al., 2021a; Zhou et al., 2021). These variants harbor mutations in the S protein that reduce neutralization by antibodies, including E484K, which is located in the RBD and is present in both, B.1.351 and P1 (Li et al., 2021; Liu et al., 2021; Wang et al., 2021c). At present, evasion form antibodies is most prominent for variant B.1.351 but it is unclear whether variants can arise that exhibit increased or even complete neutralization resistance.

India has seen a steep increase in COVID-19 cases and deaths in the recent weeks (W.H.O., 2021). It is believed that many cases are due to infection with a novel variant, B.1.617, that harbors eight mutations in the S protein, including mutations L452R and E484Q within the RBD, which introduce changes at amino acid positions known to modulate antibody-mediated neutralization (Li et al., 2021; Li et al., 2020). However, it is at present unknown whether B.1.617 evades antibody-mediated neutralization. Similarly, it is unknown whether the variant exhibits an altered dependence on host cell factors for entry, which may alter cell tropism, entry efficiency and sensitivity to entry inhibitors.

Here, we report that the S protein of B.1.617 mediates moderately enhanced entry into the human lung- and intestine-derived cell lines Calu-3 and Caco-2, respectively, and that entry is inhibited by soluble ACE2 and Camostat, the latter of which targets TMPRSS2. In contrast, entry driven by the B.1.617 S protein was fully resistant to neutralization by a monoclonal antibody with EUA for COVID-19 treatment (Bamlanivimab). Finally, B.1.617 S protein-driven entry was partially resistant against neutralization by antibodies elicited upon infection or vaccination with the Comirnaty/BNT162b2 vaccine.

## RESULTS

### The S protein of variant B.1.617 mediates increased entry into certain human intestinal and lung cell lines

The S protein of SARS-CoV-2 variant B.1.617 (GISAID Accession ID: PI_ISL_1360382) harbors a total of eight mutations compared to the S proteins of the viruses sampled at the start of the pandemic (Figure 1). Seven mutations are located within the S1 subunit, three of which are present in the N-terminal domain (R21T, E154K, Q218H), two in the RBD (L452R, E484Q) and two between the RBD and the S1/S2 border (D614G, P681R). One additional mutation is located within the S2 subunit (H1101D) (Figure 1). Since the rapid spread of variant B.1.617 in India might be due to an increased ability to enter cells or to infect a broader range of target cells, we analyzed the cell tropism and entry efficiency of variant B.1.617. For this, we employed vesicular stomatitis virus (VSV) particles pseudotyped with the S protein of either wildtype (WT) SARS-CoV-2 (Wuhan-1 isolate with D614G exchange) or variants B.1.617 or B.1.351. These pseudotyped particles faithfully mimic cell entry of SARS-CoV-2 and have been previously used to identify host factors required for SARS-CoV-2 cell entry and to study neutralization of SARS-CoV-2 by antibodies (Hoffmann et al., 2021a; Hoffmann et al., 2020; Riepler et al., 2020; Schmidt et al., 2020).

**Figure 1.**
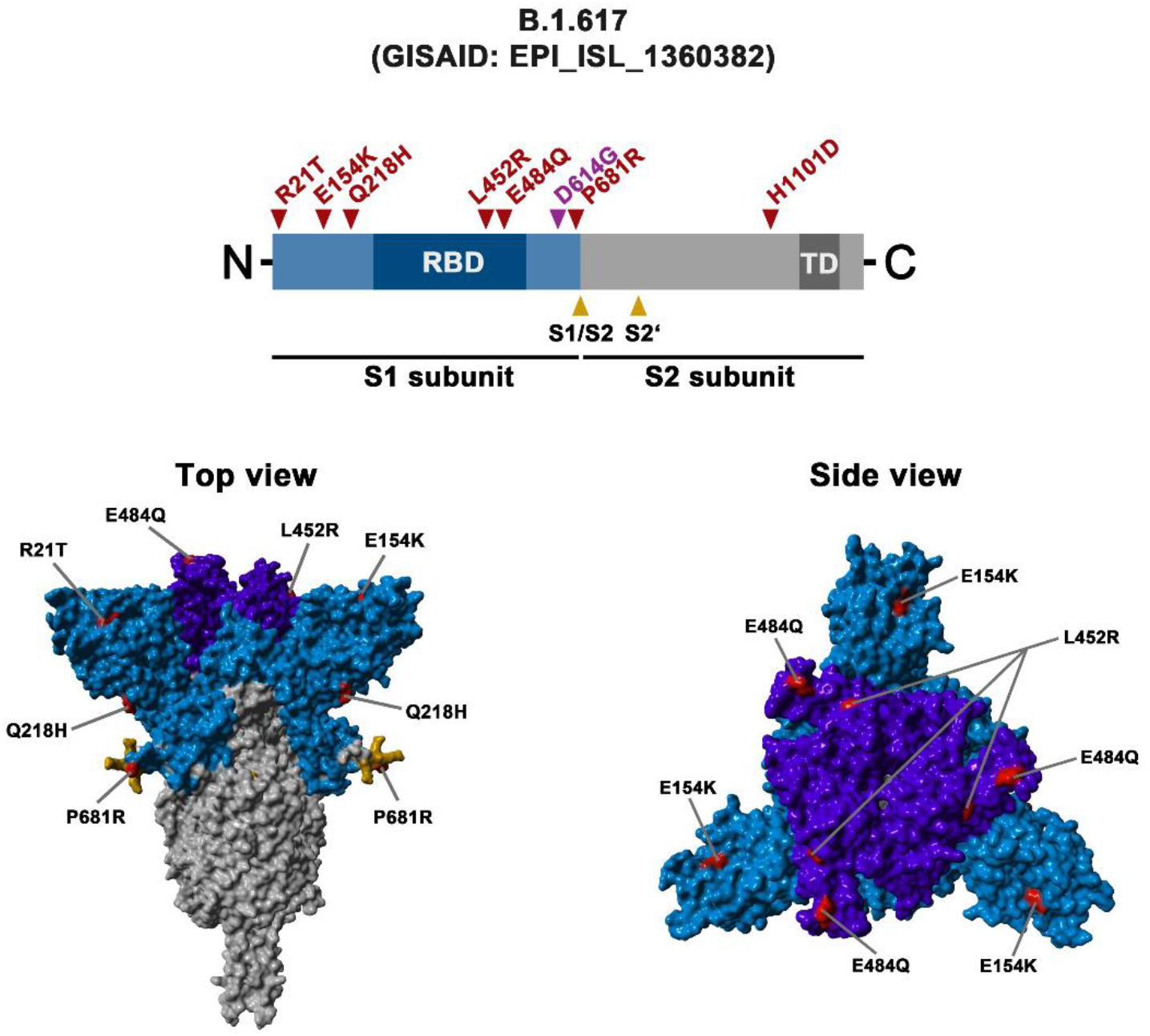
Schematic overview of the S protein from SARS-CoV-2 variant B.1.617. The location of the mutations in the context of the B.1.617 S protein domain organization is shown in the upper panel. RBD, receptor-binding domain; TD, transmembrane domain. The location of the mutations in the context of the trimeric S protein is shown in the lower panels. Color code: light blue, S1 subunit with RBD in dark blue; gray, S2 subunit; orange, S1/S2 and S2’ cleavage sites; red, mutated amino acid residues.

We analyzed a total of eight cell lines - Vero, Caco-2, Calu-3, Calu-3 (ACE2), 293T, A549 (ACE2), A549 (ACE2+TMPRSS2) and Huh-7 - most of which are commonly used as cell culture models for various aspects of SARS-CoV-2 replication. Calu-3 (ACE2), A549 (ACE2) and A549 (ACE2+TMPRSS2) cells were engineered to express ACE2 or ACE2 in conjunction with TMPRSS2.

For most cell lines we did not observe significant differences in cell entry efficiency between the WT and variant S proteins (Figure 2A and Figure S1). The only exceptions were Caco-2 and Calu-3 cells (and 293T cells in case of B.1.351), which are derived from human intestine and lung, respectively, and for which the S protein of B.1.617 (and B.1.351) mediated entry with moderately increased efficiency (Figure 2A and Figure S1). Of note this increase was less prominent when Calu-3 cells were engineered to overexpress ACE2. In addition, we investigated S protein-driven cell entry into BHK-21 cells, which were transfected either with empty plasmid or ACE2 expression plasmid. As expected, BHK-21 cells transfected with empty plasmid did not allow entry driven by any of the S proteins tested but were efficiently entered by particles bearing the VSV glycoprotein (VSV-G) (Figure 2B). In contrast, directed ACE2 expression rendered these cells susceptible to S protein-driven entry and entry efficiency was comparable between WT, B.1.351 and B.1.617 S proteins (Figure 2B). In sum, we demonstrate that the S protein of SARS-CoV-2 B.1.617 allows for moderately enhanced entry into certain cells of the respiratory and digestive tracts.

**Figure 2.**
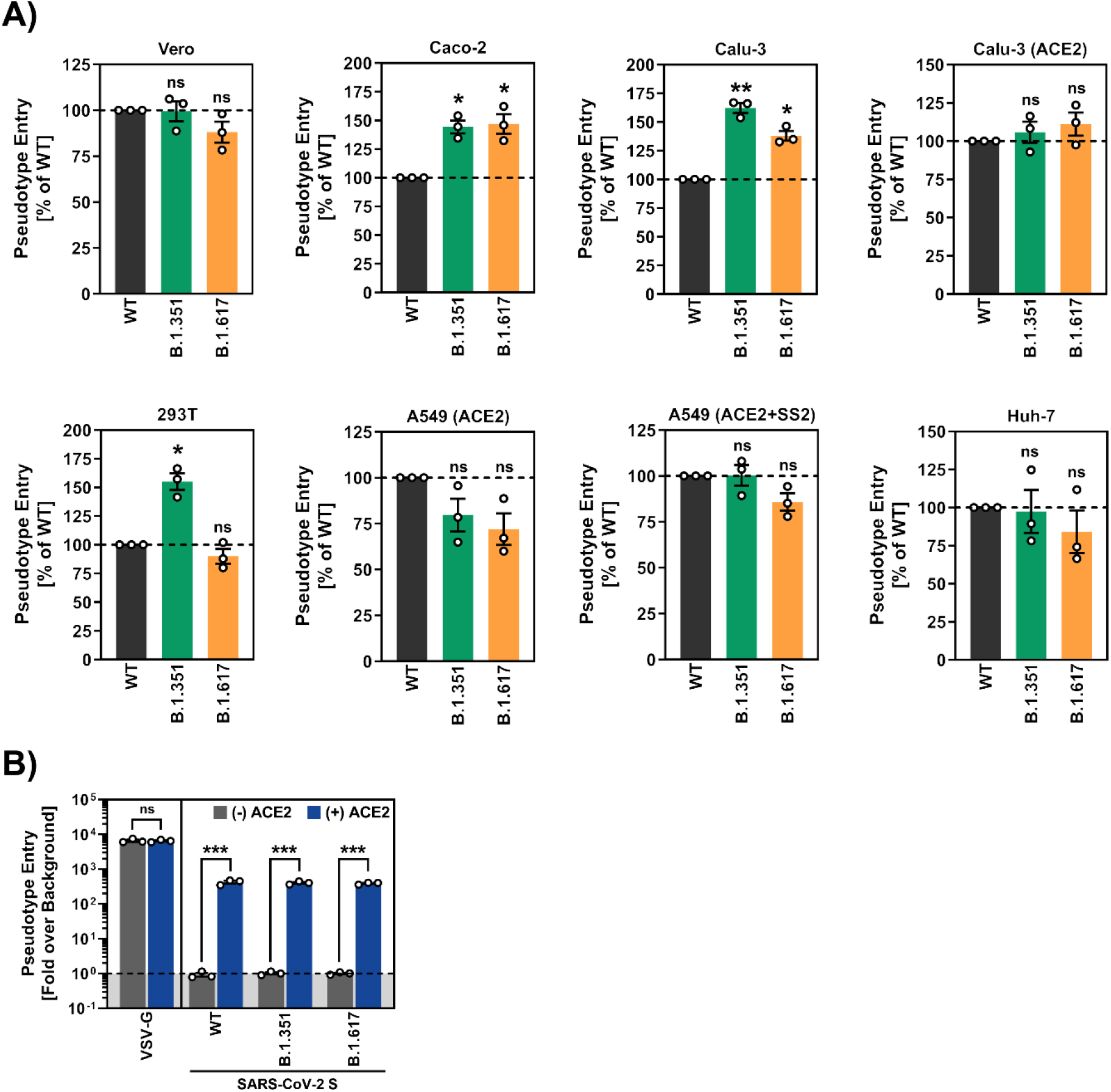
The S protein of SARS-CoV-2 variant B.1.617 S drives efficient entry into human cell lines. (A) The S protein of the SARS-CoV-2 variant B.1.617 mediates robust entry into cell lines. The indicated cell lines were inoculated with pseudotyped particles bearing the S proteins of the indicated SARS-CoV-2 variants or wild-type (WT) SARS-CoV-2 S. Transduction efficiency was quantified by measuring virus-encoded luciferase activity in cell lysates at 16-18 h post transduction. Presented are the average (mean) data from three biological replicates (each conducted with technical quadruplicates) for which transduction was normalized against SARS-CoV-2 S WT (= 100%). Error bars indicate the standard error of the mean (SEM). Statistical significance of differences between WT and variant S proteins was analyzed by one-way analysis of variance (ANOVA) with Dunnett’s posttest (p > 0.05, not significant [ns]; p ≤ 0.05, *; p ≤ 0.01, **; p ≤ 0.001, ***). See also Figure S1. (B) BHK-21 cells transfected with empty plasmid or ACE2 plasmid were inoculated with pseudotyped particles bearing the indicated S proteins, VSV-G or no viral glycoprotein (control, not shown). Presented are the average (mean) data from three biological replicates (each conducted with technical quadruplicates) for which transduction was normalized against the background (signal obtained for particles without viral glycoprotein, = 1, indicated by the dashed line). Error bars indicate the SEM. Statistical significance of differences between empty vector and ACE2-transfected cells was analyzed by multiple t-test with correction multiple comparison (Holm-Sidak method; p > 0.05, ns; p ≤ 0.001, ***).

### Soluble ACE2 and Camostat inhibit cell entry driven by the S protein of variant B.1.617

We next examined whether entry of B.1.617 can be blocked by inhibitors targeting the RBD (soluble ACE2) and proteolytic activation (Camostat) of the S protein. Soluble ACE2 binds to the RBD and blocks subsequent engagement of membrane bound ACE2. It inhibits SARS-CoV and SARS-CoV-2 cell entry and is being developed for COVID-19 treatment (Kuba et al., 2005; Monteil et al., 2020). Soluble ACE2 blocked Caco-2 cell entry driven by WT, B.1.351 and B.1.617 S proteins with comparable efficiency but did not interfere with entry driven by VSV-G (Figure 3A). Similar results were obtained for Camostat, a clinically-proven serine protease inhibitor active against TMPRSS2 (Hoffmann et al., 2020) (Figure 3B). These results indicate that soluble ACE2 and Camostat will be active against the B.1617 variant.

**Figure 3.**
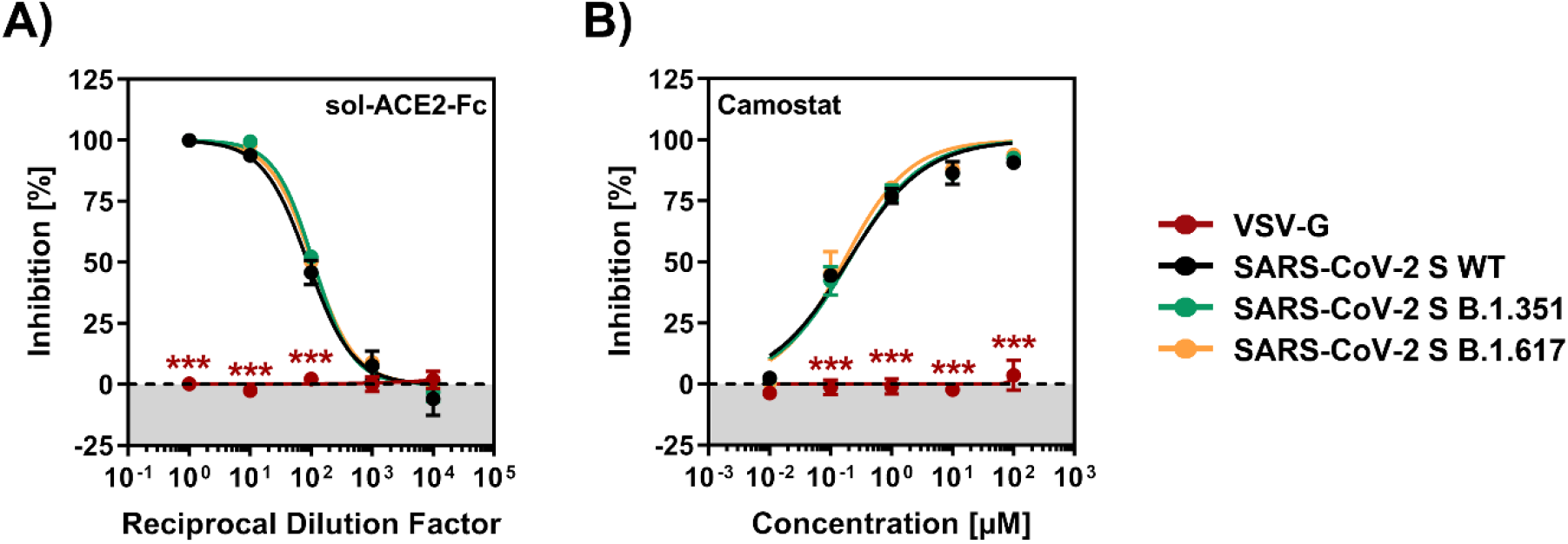
Entry driven by the S protein of SARS-CoV-2 variant B.1.617 can be blocked with soluble ACE2 and Camostat mesylate. (A) S protein-bearing particles were incubated with different concentrations of soluble ACE2 (sol-ACE2) for 30 min at 37 °C, before the mixtures were inoculated onto Caco-2 cells. (B) Caco-2 target cells were pre-incubated with different concentrations of the serine protease inhibitor Camostat mesylate for 1 h at 37 °C, before S protein-bearing particles were added. All panels: Transduction efficiency was quantified by measuring virus-encoded luciferase activity in cell lysates at 16-18 h post-transduction. For normalization, SARS-CoV-2 S protein-driven entry in the absence of sol-ACE2 or Camostat was set as 0% inhibition. Presented are the average (mean) data from three biological replicates (each performed with technical quadruplicates. Error bars indicate the SEM. Statistical significance of differences between WT and variant S proteins or VSV-G was analyzed by two-way ANOVA with Dunnett’s posttest (p > 0.05, ns [not indicated]; p ≤ 0.001, ***).

### Resistance against the therapeutic antibody Bamlanivimab

The neutralizing monoclonal antibodies Casirivimab (REGN10933), Imdevimab (REGN10987) and Bamlanivimab (LY-CoV555) and Etesevimab (LY-CoV016) (Figure S2) have received EUA for COVID-19 therapy. We analyzed whether these antibodies were able to inhibit host cell entry driven by the S protein of variant B.1.617. An irrelevant control antibody (hIgG) failed to block entry mediated by all viral glycoproteins tested (Figure 4), as expected. Casirivimab inhibited host cell entry mediated by the S protein of B.1.351 with reduced efficiency, in keeping with published data (Hoffmann et al., 2021a), and also inhibition of B.1.617 S protein-driven entry was diminished (Figure 4), in keeping with the presence of mutations in the antibody binding site (Figure S2). In contrast, B.1.351 and B.1.617 S protein-mediated entry was efficiently inhibited by Imdevimab and by a cocktail of Casirivimab and Imdevimab, termed REGN-COV (Figure 4). Further, Bamlanivimab failed to inhibit entry driven by the S protein of variant B.1.351, as expected (Hoffmann et al., 2021a), and was also unable to block entry driven by the S protein of variant B.1.617 (Figure 4). Bamlanivimab resistance of B.1.617 is in agreement with mutations in the epitope recognized by the antibody (Figure S2). Etesevimab blocked entry driven by WT and B.1.617 S proteins with comparable efficiency but failed to inhibit B.1.351 S protein-driven cell entry (Figure 4). Finally, a cocktail of Bamlanivimab and Etesevimab was less effective in inhibiting B.1.617 S protein-driven cell entry compared to WT SARS-CoV-2 S and completely failed to block entry driven by B.1.351 S (Figure 4). These results suggest that Casirivimab and particularly Bamlanivimab monotherapy may not be suitable for treatment of patients infected with variant B1.617.

**Figure 4.**
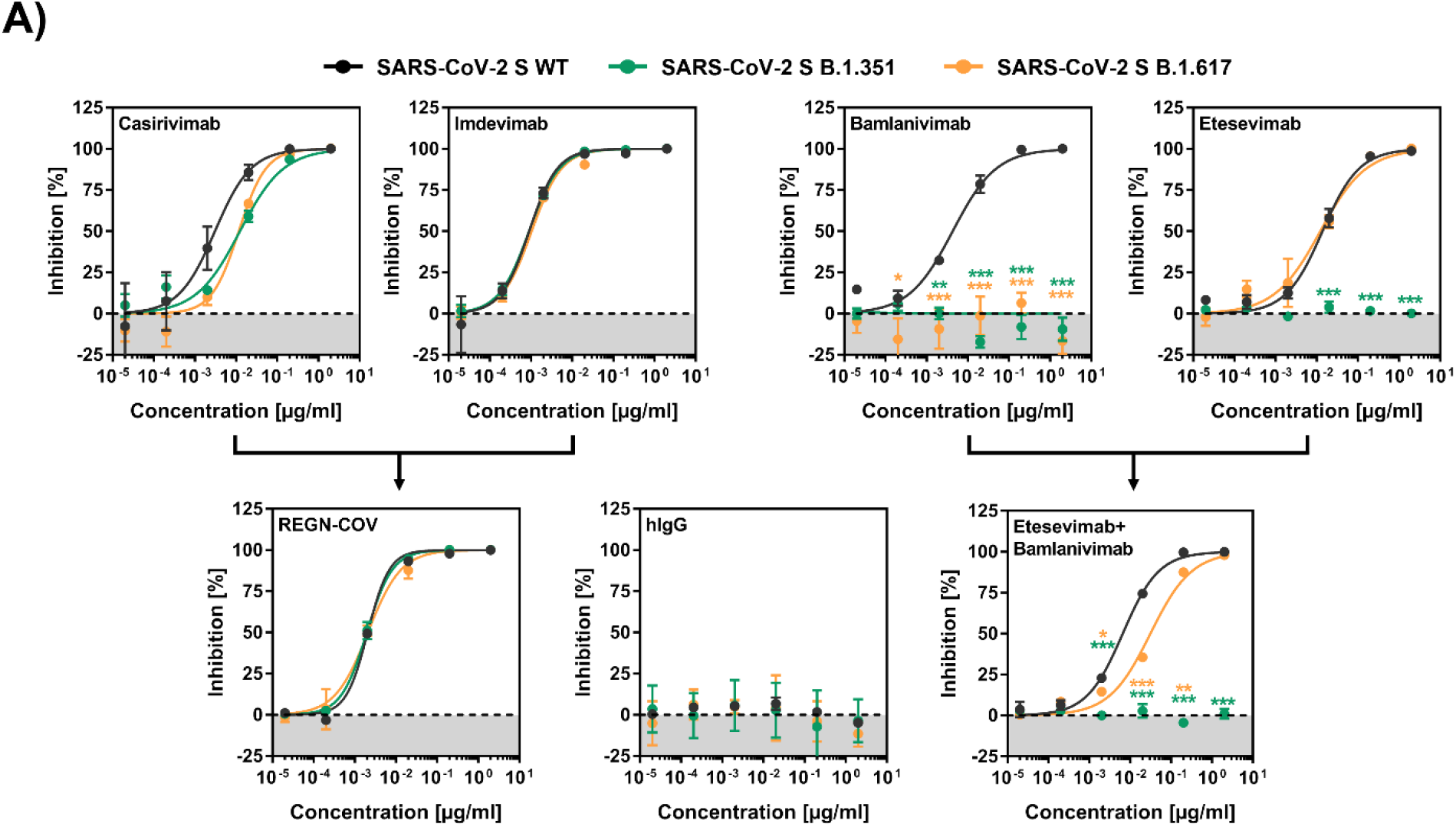
The S protein of SARS-CoV-2 variant B.1.617 is resistant to neutralization by Bamlanivimab. S protein-bearing particles were incubated with different concentrations of monoclonal antibodies for 30 min at 37 °C before the mixtures were inoculated onto Vero cells. Transduction efficiency was quantified by measuring virus-encoded luciferase activity in cell lysates at 16-18 h posttransduction. For normalization, SARS-CoV-2 S protein-driven entry in the absence of monoclonal antibody was set as 0% inhibition. Presented are the average (mean) data from three biological replicates (each performed with technical quadruplicates). Error bars indicate the SEM. Statistical significance of differences between WT and variant S proteins was analyzed by twoway ANOVA with Dunnett’s posttest (p > 0.05, ns [not indicated]; p ≤ 0.05, *; p ≤ 0.01, **; p ≤ 0.001, ***). See also Figure S2.

### Diminished neutralization by plasma from COVID-19 convalescent patients

SARS-CoV-2 infection induces the generation of neutralizing antibodies in most infected patients and it is believed that these antibody responses are important for protection from re-infection (Rodda et al., 2021; Wajnberg et al., 2020). Therefore, we determined whether variant B.1.617 evades inhibition by antibodies, which might contribute to its increasing transmission dynamics. For this, we analyzed antibody-mediated neutralization using plasma samples obtained from 15 COVID-19 patients at the intensive care unit of Göttingen University Hospital (Table S1). These plasma samples were prescreened for neutralizing activity and tested for their ability to block host cell entry driven by WT S protein and the S protein of variant B.1.617. The S protein of variant B.1.351 served as control since this S protein efficiently evades antibody-mediated neutralization (Hoffmann et al., 2021a). Inhibition of entry driven by the B.1.351 S protein was almost 6-fold less efficient as compared to WT S protein (Figure 5A and Figure S3A). Neutralization of particles bearing the S protein of variant B.1.617 was also reduced but the reduction was less prominent (~ 2-fold) (Figure 5A and Figure S3A). These results suggest that variant B.1.617 might evade antibody-mediated control in COVID-19 convalescent patients, although with moderate efficiency.

**Figure 5.**
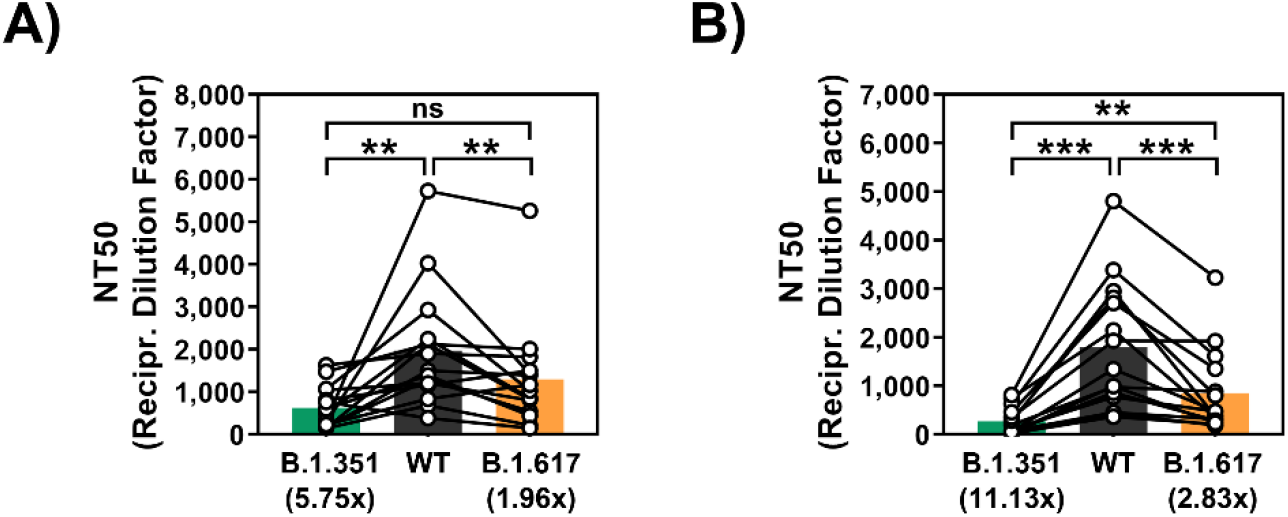
The S protein of SARS-CoV-2 variant B.1.617 evades neutralization by antibodies induced upon infection or vaccination with BNT162b2. The S protein of SARS-CoV-2 variant B.1.617 evades neutralization by convalescent plasma (A) or plasma from BNT162b2-vaccinated individuals (B). Both panels: S protein-bearing particles were incubated with different plasma dilutions (derived from infected or vaccinated individuals) for 30 min at 37 °C before the mixtures were inoculated onto Vero cells. Transduction efficiency was quantified by measuring virus-encoded luciferase activity in cell lysates at 16-18 h post-transduction and used to calculate the plasma/serum dilution factor that leads to 50 % reduction in S protein-driven cell entry (neutralizing titer 50, NT50). Presented are the average (mean) data from a single biological replicate (performed with technical quadruplicates). Error bars indicate the standard deviation. Identical plasma samples are connected with lines and the numbers in brackets indicate the average (mean) reduction in neutralization sensitivity for the S proteins of the respective SARS-CoV-2 variants. Statistical significance of differences between WT and variant S proteins was analyzed by paired two-tailed Students t-test (p > 0.05, ns; p ≤ 0.01, **; p ≤ 0.001, ***). See also Figure S3.

### Diminished neutralization by plasma from Comirnaty/BNT162b2 vaccinated patients

Vaccination with Comirnaty/BNT162b2 has been shown to be safe and to protect against COVID-19 with high efficiency (Polack et al., 2020). The vaccine is based on an mRNA that encodes the SARS-CoV-2 S protein. The vaccine induces antibody and T cell responses (Grifoni et al., 2020; Peng et al., 2020) and the neutralizing antibodies triggered by vaccination are believed to be important for vaccine-induced protection against SARS-CoV-2 infection.

Therefore, we analyzed whether cell entry driven by the S protein of variant B.1.617 can be efficiently inhibited by plasma from Comirnaty/BNT162b2 vaccinated individuals (Table S2). To address this question, we analyzed neutralization by 15 plasma samples obtained from vaccinees two to three weeks after they had received the second vaccine dose. All sera efficiently inhibited entry driven by WT S protein (Figure 5B and Figure S3B). Inhibition of entry driven by B.1.351 S protein was more than 11-fold reduced as compared to WT (Figure 5B), in keeping with expectations (Hoffmann et al., 2021a). Entry mediated by the S protein of variant B.1.617 was also less efficiently inhibited but evasion from antibody-mediated inhibition was less pronounced (~ 3-fold reduction) (Figure 5B). Thus, variant B.1.617 can partially evade control by antibodies induced by vaccination with Comirnaty/BNT162b2.

## DISCUSSION

The recent surge in COVID-19 cases and deaths in India is paralleled by the spread of the novel SARS-CoV-2 variant B.1.617. This variant harbors mutations in the RBD and other parts of the S protein that might alter important biological properties of the virus, including the efficiency of entry into target cells and the susceptibility to drugs and antibodies targeting the entry process. The present study reveals that the B.1.617 S protein can facilitate entry into Calu-3 lung and Caco-2 colon cells with slightly increased efficiency and shows that entry can be blocked by soluble ACE2 and Camostat. In contrast, Bamlanivimab, a recombinant antibody with EUA did not inhibit entry driven by the B.1.617 S protein and evidence for moderate evasion of antibodies induced by infection and Comirnaty/BNT162b2 vaccination was obtained.

The spread of variant B.1.617 in India might be partially due to increased fitness of this variant, i.e. an increased ability to amplify in patients and to be transmitted between patients. Augmented fitness has been documented for SARS-CoV-2 variant B.1.1.7 (Frampton et al., 2021; Graham et al., 2021; Leung et al., 2021) but the underlying mechanism is unclear. Increased host cell entry due to more efficient ACE2 binding or more robust proteolytic activation of the S protein might contribute to augmented fitness. Host cell entry driven by the B.1.617 S protein was not increased when cell lines were examined that expressed moderate levels of endogenous ACE2, like 293T cells, arguing against more efficient ACE2 usage by variant B.1.617. Similarly, the comparable inhibition of particles bearing WT or B.1.617 S protein by soluble ACE2 points towards no substantial differences in ACE2 binding efficiency. However, the B.1.617 S protein facilitated moderately increased entry into the human lung and intestinal cell lines Calu-3 and Caco-2, respectively, and this effect was much less pronounced when ACE2 was overexpressed in Calu-3 cells. Therefore, one can speculate that B.1.617 may have an increased ability to use certain entry augmenting factors that are expressed in a cell type specific fashion. Potential candidates are heparan sulfate (Clausen et al., 2020), Axl (Wang et al., 2021b) and neuropilin-1 (Cantuti-Castelvetri et al., 2020).

Another explanation for the increased spread of variant B.1.617 in India might be immune evasion, i.e. the ability to spread in a population in which a substantial portion of individuals has preexisting immune responses against SARS-CoV-2. This is the case in India, at least in certain areas or resource-poor communities, in which seroprevalence can be higher than 70% (Kar et al., 2021; Malani et al., 2021; Mohanan et al., 2021). The RBD of the B.1.617 S protein harbors two mutations associated with (L452R) or suspected (E484Q) of antibody evasion. Thus, mutation L452R was previously shown to facilitate escape from neutralizing antibodies (Li et al., 2020). Moreover, E484K present the B.1.351 and P.1 variants confers antibody resistance (Li et al., 2021) and one could speculate that exchange E484Q might have a similar effect. In fact, both L452R and E484K are likely responsible for B.1.617 resistance to Bamlanivimab (Starr et al., 2021) (Figure S2). In the light of these findings it was not unexpected that evasion of the B.1.617 S protein from antibodies in plasma from COVID-19 convalescent patients and Comirnaty/BNT162b2 vaccinated individuals was observed. However, evasion from antibody-mediated neutralization by B.1.617 S was clearly less pronounced as compared to the S protein of the B.1.351 variant. Differences between these variants regarding the additional mutations located in and outside of the RBD likely account for their differential neutralization sensitivity.

Collectively, our results suggest that although B.1.617 may be able to evade control by antibodies to some extend other factors might contribute to its fast spread, including a potential fitness benefit or reduced adherence to COVID-19 protection measures (e.g. mask wearing and social distancing).

## ACKNOWLEDGMENTS

We like to thank Roberto Cattaneo, Georg Herrler, Stephan Ludwig, Andrea Maisner, Thomas Pietschmann and Gert Zimmer for providing reagents. We gratefully acknowledge the originating laboratories responsible for obtaining the specimens and the submitting laboratories where genetic sequence data were generated and shared via the GISAID Initiative, on which this research is based. J.M. acknowledges funding by a Collaborative Research Centre grant of the German Research Foundation (316249678 – SFB 1279), the European Union’s Horizon 2020 research and innovation programme under grant agreement No 101003555 (Fight-nCoV) and the Federal Ministry of Economics, Germany (Combi-CoV-2). S.P. acknowledges funding by BMBF (RAPID Consortium, 01KI1723D and 01KI2006D; RENACO, 01KI20328A, 01KI20396, COVIM consortium), the county of Lower Saxony and the German Research Foundation (DFG). N.K. acknowledges funding by BMBF (ANI-CoV, 01KI2074A). M.S.W. received unrestricted funding from Sartorius AG, Lung research.

## AUTHOR CONTRIBUTIONS

Conceptualization, M.H., S.P.; Funding acquisition, S.P.; Investigation, M.H., H.H.-W., N.K., A.K., I.N., L.G., A.S., A.-S.M.; Essential resources, M.S.W., S.S., H.-M.J., M.V.S., G.M.N.B.; Writing, M.H., S.P.; Review and editing, all authors.

## MATERIALS AND METHODS

### Cell culture

293T (human, female, kidney; ACC-635, DSMZ, RRID: CVCL_0063), Huh-7 (human, male, liver; JCRB0403, JCRB; RRID: CVCL_0336, kindly provided by Thomas Pietschmann, TWINCORE, Centre for Experimental and Clinical Infection Research, Hannover, Germany), BHK-21 (Syrian hamster, male, kidney; ATCC Cat# CCL-10; RRID: CVCL_1915, kindly provided by Georg Herrler, University of Veterinary Medicine, Hannover, Germany) and Vero76 cells (African green monkey, female, kidney; CRL-1586, ATCC; RRID: CVCL_0574, kindly provided by Andrea Maisner, Institute of Virology, Philipps University Marburg, Marburg, Germany) were cultivated in Dulbecco’s modified Eagle medium (DMEM) supplemented with 10% fetal bovine serum (FCS, Biochrom or GIBCO), 100 U/ml of penicillin and 0.1 mg/ml of streptomycin (PAN-Biotech). Caco-2 (human, male, intestine; HTB-37, ATCC, RRID:CVCL_0025), Calu-3 (human, male, lung; HTB-55, ATCC; RRID: CVCL_0609, kindly provided by Stephan Ludwig, Institute of Virology, University of Münster, Germany) and Calu-3 cells stably overexpressing ACE2, Calu-3 (ACE2) (Hoffmann et al., 2021c), were cultivated in minimum essential medium supplemented with 10% FCS, 100 U/ml of penicillin and 0.1 mg/ml of streptomycin (PAN-Biotech), 1x non-essential amino acid solution (from 100x stock, PAA) and 1 mM sodium pyruvate (Thermo Fisher Scientific). Calu-3 (ACE2) cells further received 0.5 μg/ml puromycin (Invivogen). A549-ACE2 (Hoffmann et al., 2021a) and A549-ACE2/TMPRSS2 cells (Hoffmann et al., 2021a) were derived from parental A549 cells (human, male, lung; CRM-CCL-185, ATCC, RRID:CVCL_0023; kindly provided by Georg Herrler) and cultivated in DMEM/F-12 Medium (ThermoFisher Scientific) supplemented with 10% FCS, 100 U/ml of penicillin and 0.1 mg/ml of streptomycin (PAN-Biotech), 1x non-essential amino acid solution (from 100x stock, PAA), 1 mM sodium pyruvate (Thermo Fisher Scientific) and 0.5 μg/ml puromycin (Invivogen). A549-ACE2/TMPRSS2 cells further received 1 μg/ml blasticidin (Invivogen).

All cell lines were incubated at 37 °C in a humidified atmosphere containing 5% CO_2_. Authentication of cell lines was performed by STR-typing, amplification and sequencing of a fragment of the cytochrome c oxidase gene, microscopic examination and/or according to their growth characteristics. In addition, cell lines were routinely tested for contamination by mycoplasma.

### Expression plasmids

Expression plasmids for DsRed (Hoffmann et al., 2020), vesicular stomatitis virus (VSV, serotype Indiana) glycoprotein (VSV-G) (Brinkmann et al., 2017), WT SARS-CoV-2 S (codon-optimized, based on the Wuhan/Hu-1/2019 isolate, contains D614G exchange; with a C-terminal truncation of 18 amino acid) (Hoffmann et al., 2021a), SARS-CoV-2 S B.1.351 (codon-optimized; with a C-terminal truncation of 18 amino acid) (Hoffmann et al., 2021a), angiotensin-converting enzyme 2 (ACE2) (Hoffmann et al., 2013) and soluble ACE2 (Hoffmann et al., 2021a) have been described before. In order to generate the expression vector for the S protein of SARS-CoV-2 variant B.1.617, the required mutations were inserted into the WT SARS-CoV-2 S sequence by overlap extension PCR. The resulting open reading frame was further inserted into the pCG1 plasmid (kindly provided by Roberto Cattaneo, Mayo Clinic College of Medicine, Rochester, MN, USA), making use of the unique BamHI and XbaI restriction sites. Sequence integrity was verified by sequencing using a commercial sequencing service (Microsynth SeqLab). Specific cloning details (e.g., primer sequences and restriction sites) are available upon request. Transfection of 293T cells was carried out by the calcium-phosphate precipitation method, while BHK-21 cells were transfected using Lipofectamine LTX (Thermo Fisher Scientific).

### Sequence analysis and protein models

The S protein sequence of SARS-CoV-2 S variant B.1.617 (GISAID Accession ID: PI_ISL_1360382) was obtained from the GISAID (global initiative on sharing all influenza data) database (https://www.gisaid.org/). Protein models were generated using the YASARA software (http://www.yasara.org/index.html) and are based on PDB: 6XDG (Hansen et al., 2020),PDB: 7L3N (Jones et al., 2020) or PDB: 7C01 (Shi et al., 2020), or a template that was constructed by modelling the SARS-2-S sequence on PDB: 6XR8 (Cai et al., 2020), using the SWISS-MODEL online tool (https://swissmodel.expasy.org/).

### Preparation of vesicular stomatitis virus pseudotypes

For this study, we employed rhabdoviral pseudotype particles that are based on a replication-deficient VSV vector that lacks the genetic information for VSV-G and instead codes for two reporter proteins, enhanced green fluorescent protein and firefly luciferase (FLuc), VSV*ΔG-FLuc (kindly provided by Gert Zimmer, Institute of Virology and Immunology, Mittelhäusern, Switzerland) (Berger Rentsch and Zimmer, 2011). Pseudotyping of VSV*ΔG-FLuc was carried out according to a published protocol (Kleine-Weber et al., 2019). First, 293T cells that expressed the respective S protein, VSV-G (or no viral protein, control) following transfection were inoculated with VSV*ΔG-FLuc at a multiplicity of infection of three and incubated for 1 h at 37 °C. Next, the inoculum was aspirated and cells were washed with phosphate-buffered saline (PBS). Thereafter, cells received culture medium containing anti-VSV-G antibody (culture supernatant from I1-hybridoma cells; ATCC no. CRL-2700; except for cells expressing VSV-G, which received only medium) and incubated for 16-18 h. Then, pseudotype particles were harvested. For this, the culture supernatant was collected and centrifuged (2,000 x g, 10 min, room temperature) in order to pellet cellular debris. Finally, the clarified supernatant was aliquoted and stored at −80 °C.

### Production of soluble ACE2

For the production of soluble ACE2 fused to the Fc portion of human immunoglobulin G (IgG), sol-ACE2, 293T cells were grown in a T-75 flask and transfected with 20 μg of sol-ACE2 expression plasmid. The medium was exchanged at 10 h posttransfection and cells were further incubated for 38 h. Then, the culture supernatant was collected and the cells received fresh medium and were further incubated. The collected supernatant was further centrifuged (2,000 x g, 10 min, 4 °C) and stored at 4 °C. After an additional 24 h, the culture supernatant was harvested and centrifuged as described before. Next, the clarified supernatants from both harvests were combined, loaded onto Vivaspin protein concentrator columns with a molecular weight cut-off of 30 kDa (Sartorius) and centrifuged at 4,000 x g at 4 °C until a concentration factor of 20 was achieved. Finally, the concentrated sol-ACE2 was aliquoted and stored at −80 °C.

### Collection of serum and plasma samples

All plasma samples were heat-inactivated (56 °C, 30 min) before analysis. Further, all plasma were pre-screened for the presence of neutralizing activity against WT SARS-CoV-2 S using a pseudotype neutralization test. Convalescent plasma samples were collected from COVID-19 patients treated at the intensive care unit of the University Medicine Göttingen under approval given by the ethic committee of the University Medicine Göttingen (SeptImmun Study 25/4/19 Ü). For collection of convalescent plasma, Cell Preparation Tube (CPT) vacutainers with sodium citrate were used and plasma was collected as supernatant over the PBMC layer. Plasma from individuals vaccinated with BioNTech/Pfizer vaccine BNT162b2/ Comirnaty were obtained 24-31 days after the second dose. The study was approved by the Institutional Review Board of Hannover Medical School (8973_BO_K_2020). For vaccinated patients, blood was collected in S-Monovette^®^ EDTA tubes (Sarstedt).

### Transduction of target cells

All transduction experiments were carried out in 96-well format at a cell confluency of 50-80%. For experiments addressing cell tropism and entry efficiency, Vero, Caco-2, Calu-3, Calu-3 (ACE2), 293T, A549 (ACE2), A549 (ACE2+TMPRSS2) and Huh-7 target cells were inoculated with identical volumes of pseudotype preparations. BHK-21 cells were transfected 24 h prior to pseudotype inoculation with either empty plasmid or ACE2 expression plasmid (0.1 μg/well) using Lipofectamine LTX (Thermo Fisher Scientific). In order to study the antiviral activity of Camostat mesylate, Caco-2 target cells, which express endogenous TMPRSS2, were pre-incubated for 1 h with medium containing different concentrations (100, 10, 1, 0.1 or 0.01 μM) of Camostat mesylate (prepared from a 100 mM stock; Tocris) or the solvent dimethyl sulfoxide (DMSO, 1:1,000; Sigma-Aldrich) as control before being inoculated with VSV bearing S protein, VSV-G or no glycoprotein. For experiments addressing the antiviral activity of soluble ACE2, pseudotype particles bearing S protein, VSV-G or no protein were pre-incubated with different amounts of concentrated sol-ACE2 (final sol-ACE2 dilutions in the mixtures: undiluted, 1:10, 1:100, 1:1,000:10,000) or only medium for 30 min at 37 °C, before the mixtures were added onto Caco-2 cells. In all cases, transduction efficiency was determined at 16-18 h postinoculation. For this, the culture supernatant was aspirated. Next, cells were lysed in PBS containing 0.5% Triton-X-100 (Carl Roth) for 30 min at room temperature. Thereafter, cell lysates were transferred into white 96-well plates. Finally, FLuc activity was measured using a commercial substrate (Beetle-Juice, PJK; Luciferase Assay System, Promega) and Hidex Sense Plate Reader (Hidex).

### Pseudotype particle neutralization test

For neutralization experiments, S protein bearing pseudotype particles were pre-incubated for 30 min at 37 °C with different concentrations of monoclonal antibodies (Casirivimab, Imdevimab, Casirivimab+Imdevimab [1:1], Bamlanivimab, Esetevimab, Bamlanivimab+Etesevimab [1:1] or unrelated control IgG [2, 0.2, 0.02, 0.002, 0.0002, 0.00002 μg/ml]) or plasma samples obtained from convalescent COVID-19 patients or individuals vaccinated with the Pfizer/BioNTech vaccine BNT162b2/Comirnaty (1:25, 1:100, 1:400, 1:1,600, 1:6,400, 1:25,600), before the mixtures were inoculated onto Vero cells. In all cases, particles incubated only with medium served as control. Transduction efficiency was determined at 16-18 h postinoculation as described above.

### Data analysis

The results presented in this study represent average (mean) data obtained from three biological replicates, each conducted with technical quadruplicates. The only exception are the neutralization data involving plasma from infected/vaccinated individuals, which are based on a single experiment (standard in the field), conducted with technical quadruplicates. Error bars are defined as either standard deviation (SD, neutralization data involving plasma from infected/vaccinated individuals) or standard error of the mean (SEM, all other data). Data ware analyzed using Microsoft Excel (as part of the Microsoft Office software package, version 2019, Microsoft Corporation) and GraphPad Prism 8 version 8.4.3 (GraphPad Software).

Data normalization was performed in the following fashion: (i) For comparison of entry efficiency by the different S proteins, transduction was normalized against WT SARS-CoV-2 S (set as 100%). Alternatively, transduction was normalized against the background signal (luminescence measured for cells inoculated with particles bearing no viral glycoprotein; set as 1). (ii) In order to investigate inhibition of S protein-driven cell entry by sol-ACE2, Camostat mesylate, monoclonal antibodies or plasma from infected/vaccinated individuals, transduction was normalized against a reference sample (e.g., control-treated cells or pseudotypes, set as 0% inhibition).

The neutralizing titer 50 (NT50) value, which indicates the plasma dilution that causes a 50 % reduction of transduction efficiency, was calculated using a non-linear regression model (inhibitor vs. normalized response, variable slope). Statistical significance was tested by one- or two-way analysis of variance (ANOVA) with Dunnett’s post-hoc test, paired Students t-test or by multiple t-test with correction for multiple comparison (Holm-Sidak method). Only p values of 0.05 or lower were considered statistically significant (p > 0.05, not significant [ns]; p ≤ 0.05, *; p ≤ 0.01, **; p ≤ 0.001, ***). The details on the statistical test and the error bars are specified in the figure legends.

## SUPPLEMENTAL INFORMATION

**Figure S1.**
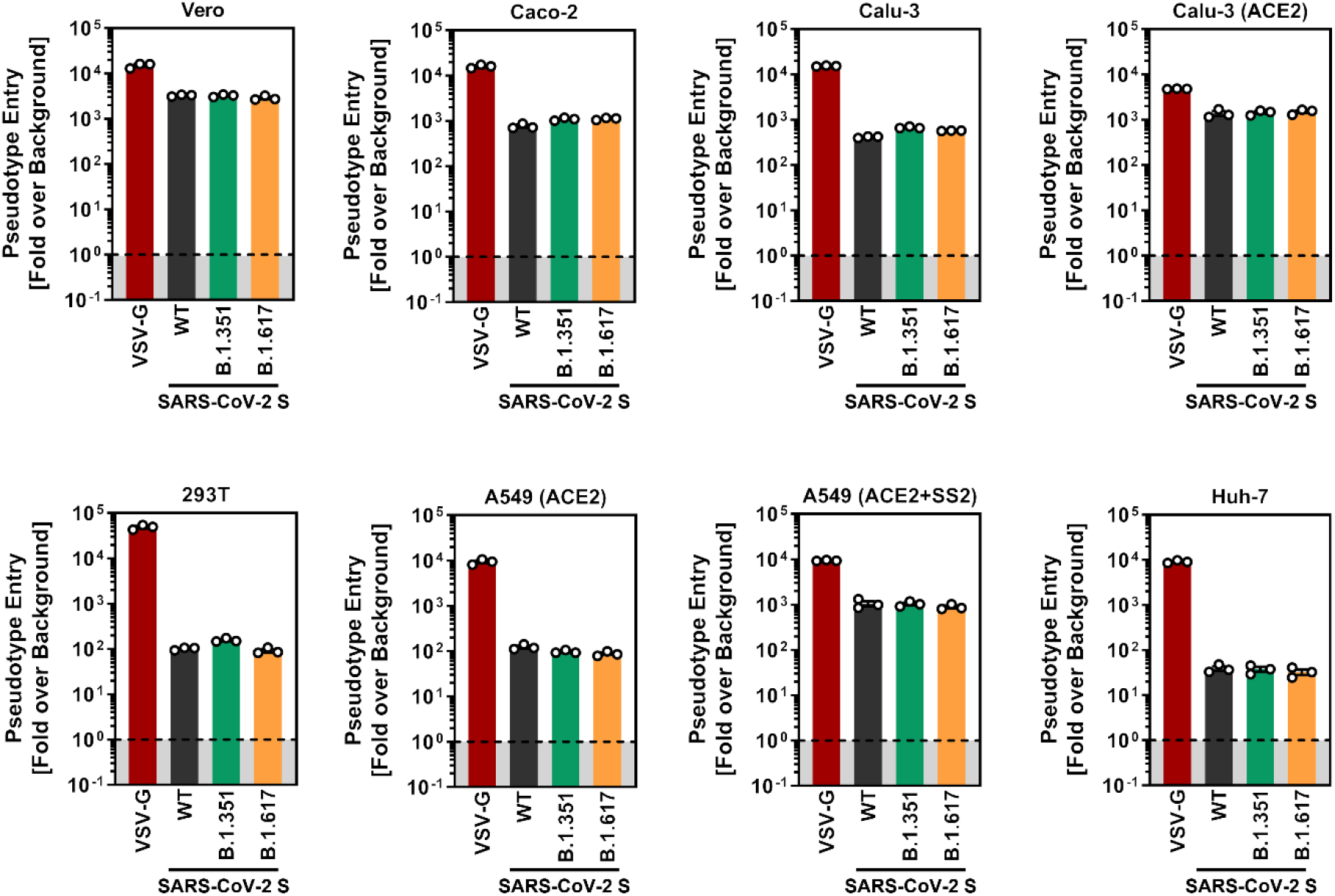
Transduction data normalized against the assay background (related to Figure 2). The experiment was performed as described in the legend of Figure 1A. Presented are the average (mean) data from the same three biological replicates (each conducted with technical quadruplicates) as presented in Figure 1A with the difference that this time transduction was normalized against signals obtained from cells inoculated with particles bearing no viral glycoprotein (background, set as 1). Further, transduction data of particles bearing VSV-G are included. Error bars indicate the SEM.

**Figure S2.**
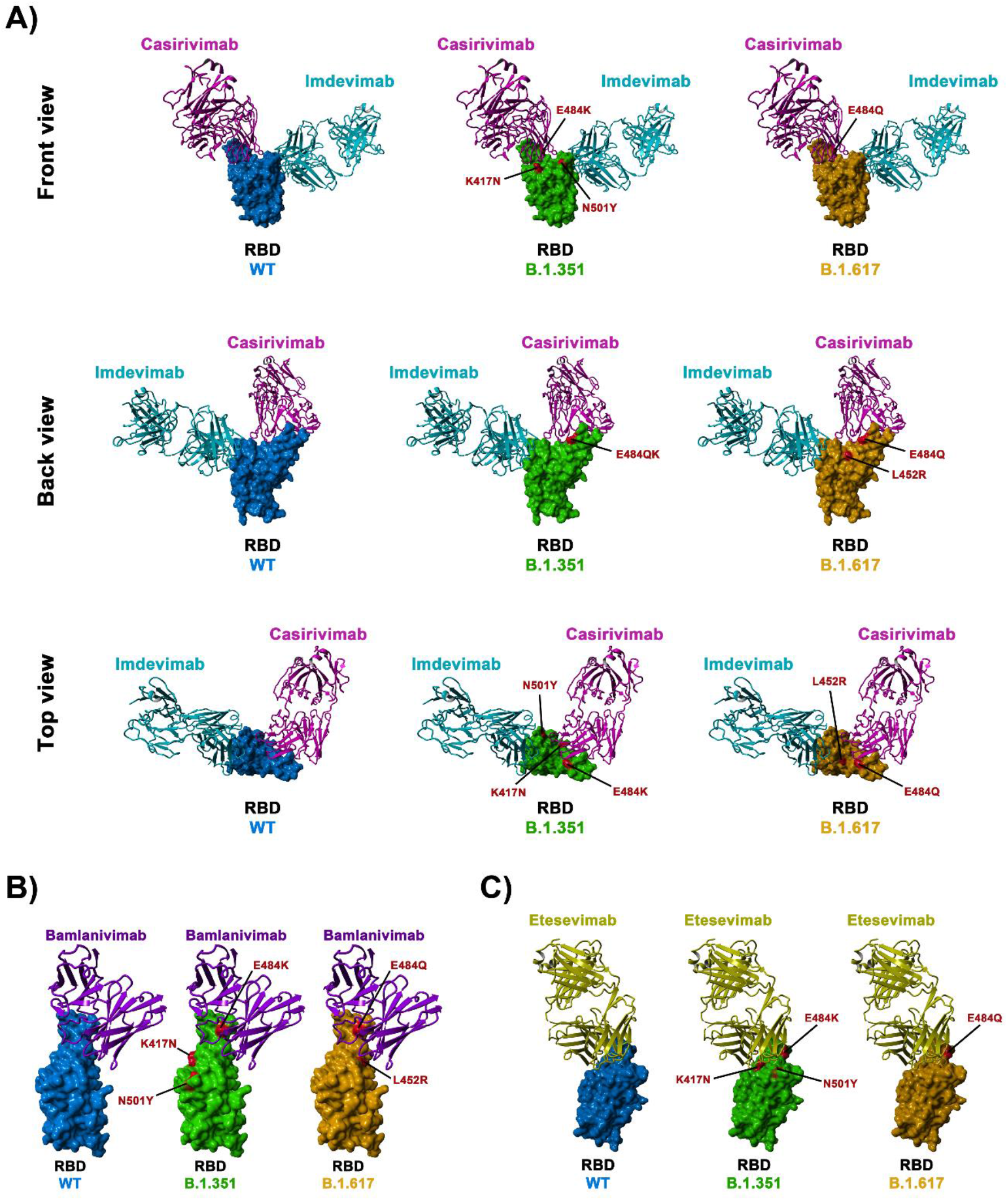
Location of SARS-2-S B.1.351 and B. 1.617 RBD mutations with respect to the binding interface of Casirivimab and Imdevimab (A), Bamlanivimab (B) and Esetevimab (C) (related to Figure 4). The protein models of the SARS-2-S receptor-binding domain (RBD, blue) in complex with antibodies Casirivimab (pink) and Imdevimab (turquoise) were constructed based on the 6XDG template (Hansen et al., 2020), while the protein models of the SARS-2-S RBD in complex with antibody Bamlanivimab (purple) and Etesevimab (yellow) were based on the PDB: 7L3N (Jones et al., 2020) and PDB: 7C01 (Shi et al., 2020) template, respectively. Residues highlighted in red indicate amino acid variations found in the SARS-CoV-2 variants.

**Figure S3.**
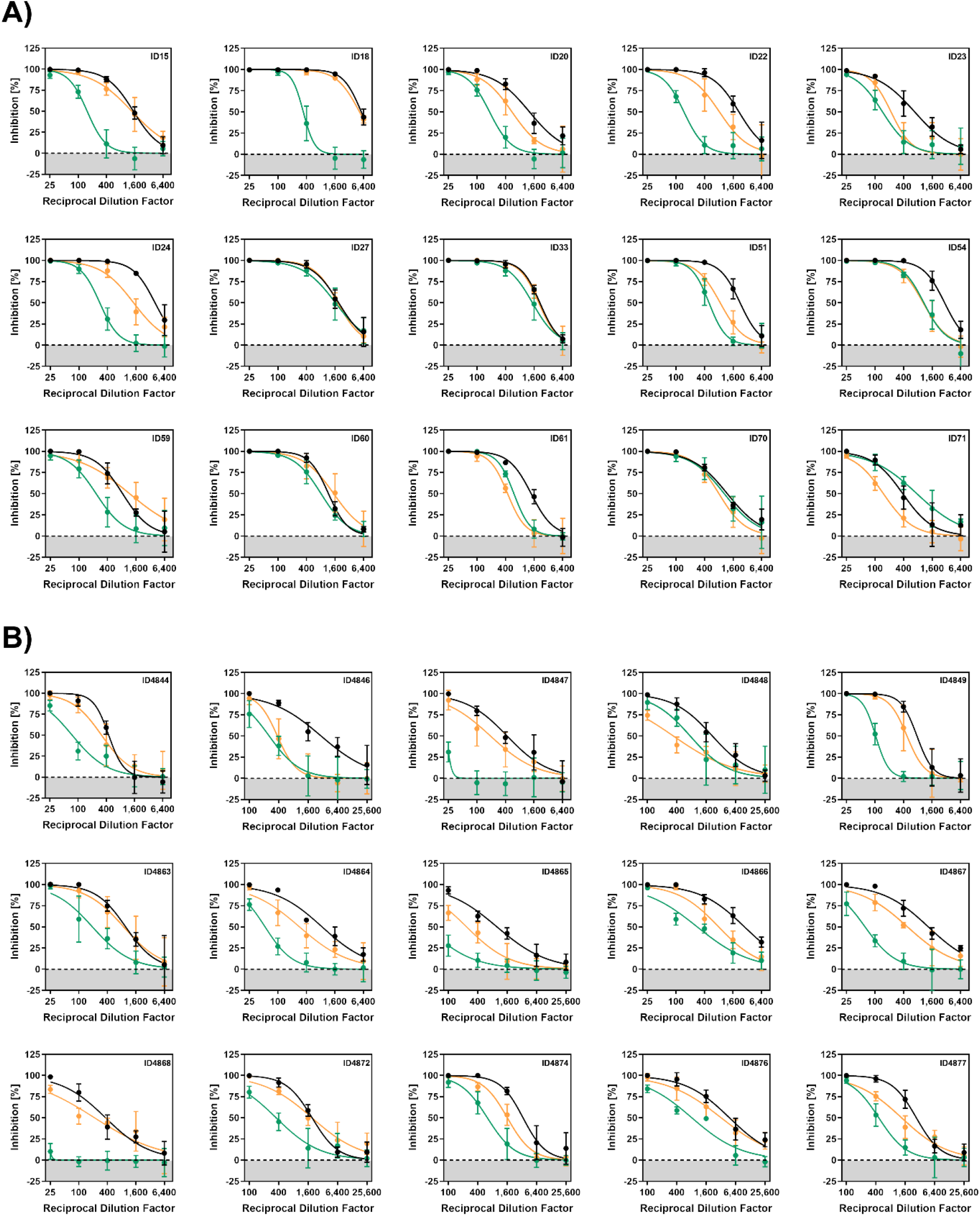
Individual neutralization data (related to Figure 5). Pseudotypes bearing the indicated S proteins were incubated (30 min, 37 °C) with different dilutions of plasma derived from COVID-19 patients (A) or individuals vaccinated with the Pfizer/BioNTech vaccine Comirnaty/BNT162b2 (B) and inoculated onto Vero target cells. Transduction efficiency was quantified by measuring virus-encoded luciferase activity in cell lysates at 16-18 h posttransduction. Presented are the data from a single representative experiment conducted with technical Quadruplicates. For normalization, inhibition of S protein-driven entry in samples without plasma was set as 0 %. Error bars indicate the SD. The data were further used to calculated the plasma/serum dilution that leads to 50% reduction in S protein-driven cell entry (neutralizing titer, NT50, shown in Figure 5).

**Table S1:** COVID-19 patient data

|  |  |  | WHO classification upon ICU admission |  |  |  |  |  |  |  |  |  |  |  |  |  |  |  |
| --- | --- | --- | --- | --- | --- | --- | --- | --- | --- | --- | --- | --- | --- | --- | --- | --- | --- | --- |
| ID | Symptoms before hospital admission (days) | Symptoms before ICU admission (days) | mild | moderate | severe | critical | Age | Sex | Diabetes | Hypertension | Cardiac disease | Chronic lung disease + Asthma | Cerebrovascular disease | Chronic kidney disease | Immunosuppression | Cancer | Obesity | Smoking |
| SI 15 | ND | ND | - | - | - | x | 65 | M |  |  | x |  |  |  |  | x |  |  |
| SI 16 | ND | ND | - | x | - | x | 71 | M |  |  |  |  |  |  |  | x |  |  |
| SI 18 | 2 | 11 | - | - | - | x | 74 | F | x | x |  |  |  |  |  | x |  |  |
| SI 20 | ND | ND | - | - | - | x | 61 | M | x |  |  | x |  |  |  |  |  |  |
| SI 22 | 5 | 5 | - | - | - | x | 25 | F |  |  |  |  |  |  |  |  | x |  |
| SI 23 | 2 | 8 | - | - | x | - | 69 | F | x |  |  |  |  |  |  |  |  |  |
| SI 24 | 4 | 8 | - | - | - | x | 61 | M |  | x |  |  |  |  |  |  |  | x |
| SI 27 | ND | ND | - | - | - | x | 52 | M | x | x |  |  |  |  |  |  |  |  |
| SI 33 | 1 | 14 | - | - | - | x | 75 | M | x | x | x |  |  |  |  |  |  |  |
| SI 51 | 4 | 12 | - | - | - | x | 71 | M |  |  |  |  |  |  |  |  |  |  |
| SI 54 | 8 | 7 | - | - | - | x | 58 | F |  | x |  | x |  |  |  | x |  |  |
| SI 59 | 6 | 3 | - | - | - | x | 46 | M |  | x | x |  |  | x |  |  |  | x |
| SI 60 | 5 | 6 | - | - | - | x | 61 | F |  |  |  | x |  |  |  |  |  |  |
| SI 61 | 7 | 2 | - | - | - | x | 50 | M |  |  |  | x |  |  |  |  | x |  |
| SI 70 | ND | ND | - | - | x | - | 34 | F |  |  |  |  |  |  |  |  |  |  |
| SI 71 | 5 | 3 | - | x | - | - | 54 | M |  | x |  |  |  |  |  |  |  |  |

ND =Not determined

**Table S2:** BNTl62b2-vaccinated patient data. BNT162b2-vaccinated patient data. Serological data shows antibody titer against spike (IgG) protein determined by quantitative ELISA (SARS-CoV-2-QuantiVac; Euroimmun, Lübeck, Germany) according to the manufacturer’s instructions. Antibody levels are expressed as RU/ml assessed from a calibration curve with values above 10 RU/mL defined as positive, values beyond the standard curve are expressed as >120 RU/ml.

| ID | Age (y) | Gender | Time since 2<sup>nd</sup> vaccination (d) | Spike-IgG |
| --- | --- | --- | --- | --- |
| 4844 | 47 | f | 31 | >120 |
| 4846 | 28 | f | 29 | >120 |
| 4847 | 57 | f | 26 | >120 |
| 4848 | 29 | m | 26 | >120 |
| 4849 | 39 | m | 24 | >120 |
| 4863 | 58 | f | 26 | >120 |
| 4864 | 53 | f | 26 | >120 |
| 4865 | 57 | m | 25 | >120 |
| 4866 | 59 | f | 29 | >120 |
| 4867 | 52 | f | 30 | 34,4 |
| 4868 | 56 | f | 30 | 119.6 |
| 4872 | 37 | f | 25 | >120 |
| 4874 | 26 | f | 29 | >120 |
| 4876 | 27 | f | 30 | >120 |
| 4877 | 29 | f | 28 | >120 |

